# AMMISOFT for AMMI Analysis with Best Practices

**DOI:** 10.1101/538454

**Authors:** Hugh G. Gauch, David R. Moran

## Abstract

The Additive Main effects and Multiplicative Interaction (AMMI) model has been used extensively for analysis of multi-environment yield trials for two main purposes: understanding complex genotype-by-environment interactions and increasing accuracy. A 2013 paper in *Crop Science* presented a protocol for AMMI analysis with best practices, which has four steps: (i) analysis of variance, (ii) model diagnosis, (iii) mega-environment delineation, and (iv) agricultural recommendations. This preprint announces free open-source software, called AMMISOFT, which makes it easy to implement this protocol and thereby to accelerate crop improvement.

---

The Additive Main effects and Multiplicative Interaction (AMMI) model has been used extensively to analyze multi-environment yield trials, which are one of the most common kinds of experiments in agricultural research. There have been several recent review articles about AMMI in *Crop Science* (Gauch, 2006; Yan et al., 2007; Gauch et al., 2008; Yang et al., 2009). AMMI software abounds in R, SAS, Genstat, CropStat, and other packages, including the senior author’s MATMODEL written in FORTRAN about 25 years ago (Gauch and Furnas, 1991).

However, all software for AMMI analysis that has come to our attention fails to implement AMMI in a manner that facilitates contemporary best practices, as described in Gauch (2013). Likewise, most published AMMI analyses fail to exhibit best practices, with the sad consequence that valuable information in costly data is wasted, thereby needlessly slowing the rate of crop improvement. A particularly flagrant problem is routine lack of model diagnosis to select the best member of the AMMI model family for a given dataset and research purpose, despite warnings that model diagnosis is essential for reliable and optimal results (Gauch et al., 2008; Yang et al., 2009).

To remedy these problems, this preprint announces the free open-source program AMMISOFT that implements AMMI analysis with best practices. AMMISOFT is user-friendly, mainly because each table and graph is accompanied by an explanation of its meaning. This software is illustrated with an international bread wheat (*Triticum aestivum* L.) yield trial (Crossa et al., 1991; Gauch, 2013).

## THE AMMI MODEL

A yield-trial experiment typically has two designs, the treatment design and the experimental design. For a number of genotypes tested in a number of environments, the treatment design is a two-way factorial of genotypes by environments. AMMI is applied to this genotypes-by-environments treatment design. Often yield trials also have an experimental design involving randomization, replication, blocking, or the use of spatial information. However, AMMISOFT can analyze only one experimental design, the randomized complete block design which is especially common. Other experimental designs need to be analyzed by other software. Of course, best practices involve statistical analysis of both designs.

AMMI first applies analysis of variance (ANOVA) to partition the variation into genotype main effects (G), environment main effects (E), and genotype-by-environment interaction effects GE), and then it applies principal components analysis (PCA) to GE. AMMI constitutes a model family because 0 or 1 or more interaction principal components (IPC) can be retained in the model before relegating higher components to a discarded residual, with these successive models being denoted by AMMI0 having no IPC, AMMI1 having 1 IPC, AMMI2 having 2 IPC, and so on, up to the full model denoted by AMMIF with expected values that equal the original data (which are the averages over replications if replicated). The AMMI model equation is:

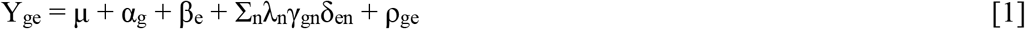

where Y_ge_ is the yield of genotype g in environment e, μ is the grand mean, α_g_ is the genotype deviation from the grand mean, β_e_ is the environment deviation, λ_n_ is the singular value for IPC n and correspondingly λ^2^_n_ is its eigenvalue, γ_gn_ is the eigenvector value for genotype g and component n, δ_en_ is the eigenvector value for environment e and component n, with both eigenvectors scaled as unit vectors, and ρ_ge_ is the residual. Commonly the interaction scores are scaled as λ^0.5^_n_γ_gn_ and λ ^0.5^ _n_δ_en_ so that their products estimate interactions directly, without the need of yet another multiplication by λ.

## PROGRAM AVAILABILITY AND EXECUTION

AMMISOFT is available for free from the senior author’s Cornell home page at https://scs.cals.cornell.edu/people/hugh-gauch. This software is open source and may be redistributed or modified under the BSD 3-Clause License published by the Regents of the University of California. To download the program, click on “Free AMMISOFT Program” at the bottom of this home page, and then “Download” at the bottom of the next page. The zipped download file occupies about 43 MB, and the unzipped files about 130 MB of memory. Installation produces an icon for AMMISOFT that has a green letter A flanked by tan heads of wheat; clicking on this icon launches the program. While AMMISOFT is running, it requires about 256 MB of high-speed memory. It has been developed for PC computers with a Windows operating system; it is not available for Apple computers. The user interface and graphs are coded in about 500 lines of Python and the AMMI computations and tables are coded in about 2,000 lines of FORTRAN. However, this software is self-contained so neither a Python interpreter nor a FORTRAN compiler are needed, unless users want to make modifications in their own versions of AMMISOFT. As explained in the instructions on this home page, installation includes a brief documentation in MS Word format about how to use AMMISOFT, and it is advisable to print this document for convenient reference.

Alternatively, AMMISOFT can be downloaded from the GitHub repository at https://github.com/tequa/ammisoft and this has the setup and all of its dependencies. Instructions on how to compile the setup are in the README file.

AMMISOFT is exceptionally user-friendly because each table and graph is accompanied by an explanation of its meaning. Consequently, merely two brief publications suffice to prepare agricultural researchers to use AMMISOFT effectively: the 10-page article on AMMI protocol (Gauch, 2013) and the 17-page program documentation included in the download.

## FOUR TABLES

AMMISOFT produces four tables in a logical order. Each is followed by an explanation of how to interpret and use the table.

The first table in AMMISOFT output is the familiar ANOVA table, as in Table 1 in the protocol paper (Gauch, 2013). If the data are replicated, the following explanation includes diagnosis of the best member of the AMMI model family for the purpose of maximizing predictive accuracy. This diagnosis is based on two considerations. First, F_R_ tests at the 0.01 significance level are used to determine how many IPCs are significant (Piepho, 1995). These tests are used, rather than those devised by Gollub (1968), because they reflect predictive accuracy much better. Second, the amount of signal and noise in the GE interactions are estimated and this provides another method for model diagnosis. These two diagnoses are compared to provide a consensus. Incidentally, the older MATMODEL software also used cross-validation for model diagnosis and it can still be used for that purpose since both programs use the same data format. However, variants of cross-validation and bootstrap methods are currently an active area of research for AMMI analysis (Forkman and Piepho, 2014). Accordingly, a decision was made to omit cross-validation, at least until a settled verdict has been reached on the best method. Also, the performance of F_R_ tests and signal-noise considerations is excellent, and their computational burden is absolutely miniscule compared to cross-validation.

**Table 1.** Genotype winners for a sequence of increasingly complex members of the Additive Main effects and Multiplicative Interaction (AMMI) model family for an international wheat trial. The simplest member, AMMI0 with no principal components, has a single winner in all 25 environments, namely the overall winner genotype 5 or KAUZ#2 (see Table 1 in Crossa et al. 1991 for the full names of the genotypes). Then AMMI1 has 3 winners, and so on, until the full model AMMIF has 12 winners, as noted at the bottom of this table. An additional 4 genotypes never win in any environment, so they are not listed. These 14 genotypes that do win in at least 1 environment are listed in order by their interaction principal component (IPC) scores. The practical choice for mega-environment delineation is AMMI1 with 3 winners or mega-environments (which can be further simplified by merging the small mega-environment with only 3 wins into the larger mega-environments). To the left of this best choice, an overly simple model would reduce yield in many environments by failing to exploit substantial narrow adaptations; whereas to the right, overly complex models would capture spurious noise that obscures winners, favors losers, and results in a totally impractical mega-environment scheme.

|  |  | AMMI Model Family |  |  |  |  |  |  |  |  |
| --- | --- | --- | --- | --- | --- | --- | --- | --- | --- | --- |
| Genotype |  | 0 | 1 | 2 | 3 | 4 | 5 | 6 | 7 | F |
| 5 | KAZ2 | 25 | 13 | 9 | 9 | 7 | 6 | 4 | 6 | 4 |
| 4 | KAZ1 |  |  | 1 | 1 | 5 | 4 | 3 | 1 | 3 |
| 7 | GENA |  |  |  |  |  | 1 | 4 | 5 | 3 |
| 9 | SERI |  | 3 | 2 | 1 | 6 | 2 | 1 |  | 1 |
| 6 | RWBB |  |  | 4 | 3 | 1 | 2 | 2 | 3 | 1 |
| 10 | BUPV |  |  |  | 1 |  |  |  |  |  |
| 18 | HDHO |  |  |  |  |  |  |  |  | 1 |
| 16 | RGKB |  |  |  |  |  | 1 | 2 | 1 | 1 |
| 17 | VEPV |  |  |  |  |  |  |  | 1 | 1 |
| 8 | AGA3 |  |  |  |  | 1 | 2 | 2 | 1 | 1 |
| 14 | BJYC |  |  |  |  |  | 1 | 1 | 2 | 4 |
| 2 | CNPR |  |  |  | 3 |  |  |  |  | 2 |
| 15 | HAPR |  | 9 | 7 | 6 | 5 | 5 | 5 | 4 | 3 |
| 11 | KETO |  |  | 2 | 1 |  | 1 | 1 | 1 |  |
| Mega-environments |  | 1 | 3 | 6 | 8 | 6 | 10 | 10 | 10 | 12 |

Table 1 is the second table in AMMISOFT output, which shows winning genotypes for the AMMI model family for the international bread wheat trial. This is a new kind of table that had not yet been invented when the protocol paper was published. Its purpose is to help with diagnosis of the best AMMI model for the purpose of mega-environment delineation. Model diagnosis depends not only on the individual yield-trial dataset, but also on the research or agricultural purpose. Hence, for a given dataset, it is possible for different AMMI models to serve different purposes, such as AMMI3 being best for predictive accuracy but AMMI1 for mega-environment delineation.

Mega-environments are distinguished by having different genotype winners. Increasingly complex AMMI models generally have more genotype winners or mega-environments, as shown in the list at the bottom of the above table, which begins with AMMI0 (the merely additive model) having 1 winner and ends with AMMIF (the full model or actual data) having 12 mega-environments. The genotypes are listed in IPC1 order, so those at the top and bottom have opposite GE interaction patterns (and another 4 genotypes never win, so they are not listed). Often this contrast in the genotypes has an evident agricultural interpretation that has a corresponding contrast in the environments (which likewise are listed in their IPC1 order in the next table). A genotype at the top of this table has positive GE interactions with environments at the top of the next table, and negative GE interactions with environments at the bottom of the next table; and the opposite patterns apply to genotypes at the bottom of this table.

Practical constraints frequently limit the number of workable mega-environments to only 2 or 3 (or perhaps a few more). This often requires a lower AMMI model than that diagnosed solely by statistical tests to optimize predictive accuracy. Fortunately, for most yield trials, this lower AMMI model is almost as predictively accurate as the best model for prediction, and is also far more accurate than AMMIF, that is, the actual data. For this wheat trial, AMMI1 is suitable, producing 3 mega-environments.

When replicated data are not available, model diagnosis must feature considerations other than statistical tests of predictive accuracy, including the agricultural interpretability of IPCs and the practicality of mega-environment schemes. Especially in this context, Table 1 can be helpful for selecting the most suitable member of the AMMI model family.

Table 2 is the third table in AMMISOFT output, which shows genotype rankings for the AMMI1 and AMMIF models, as in Table 2 in the protocol paper (except that the AMMI3 ranks in the middle of that table are not included). The environments are listed in IPC1 order, so those at the top and bottom have opposite GE interaction patterns.

**Table 2.** Ranking table showing the top 5 genotypes for 2 members of the Additive Main effects and Multiplicative Interaction (AMMI) model family. AMMI1 facilitates mega-environment delineation, and AMMIF represents the raw data. The 25 environments are ordered by their IPC1 scores, which reflect an ecological gradient from short growing seasons at the top to long growing seasons at the bottom. Blank lines separate the 3 mega-environments delineated by AMMI1, which can be simplified with little loss of yield by transferring environments 17 and 20 to the top mega-environment and environment 3 to the bottom mega-environment. Genotypes near the top of Table 1 have positive GE interactions in environments near the top of this table and negative interactions in environments near the bottom, whereas genotypes near the bottom of Table 1 have the opposite pattern. Genotype 5 is the yield trial’s overall winner, and ratio expresses the yield advantage for switching to other genotypes to exploit narrow adaptations, as explained more fully in the text. AMMI1 has few and workable mega-environments, whereas noise in the raw data AMMIF causes spurious complexity that results in 12 winners or mega-environments, which is wholly impractical.

| Environment | Ratio | AMMI1 Ranks |  |  |  |  | AMMIF Ranks |  |  |  |  |
| --- | --- | --- | --- | --- | --- | --- | --- | --- | --- | --- | --- |
|  |  | 1 | 2 | 3 | 4 | 5 | 1 | 2 | 3 | 4 | 5 |
| 25 MM | 1.0000 | 5 | 4 | 7 | 9 | 6 | 5 | 1 | 4 | 9 | 17 |
| 8 MG | 1.0000 | 5 | 4 | 7 | 9 | 6 | 5 | 4 | 18 | 12 | 15 |
| 21 TC | 1.0000 | 5 | 4 | 7 | 9 | 6 | 9 | 16 | 5 | 7 | 4 |
| 18 SC | 1.0000 | 5 | 4 | 9 | 7 | 6 | 7 | 4 | 5 | 2 | 17 |
| 12 SJ | 1.0000 | 5 | 4 | 9 | 7 | 6 | 18 | 7 | 6 | 5 | 12 |
| 15 SS | 1.0000 | 5 | 4 | 9 | 7 | 6 | 5 | 9 | 4 | 1 | 7 |
| 7 IL | 1.0000 | 5 | 4 | 9 | 7 | 6 | 6 | 9 | 16 | 4 | 5 |
| 22 PA | 1.0000 | 5 | 4 | 9 | 7 | 6 | 7 | 1 | 10 | 12 | 4 |
| 6 BJ | 1.0000 | 5 | 4 | 9 | 7 | 6 | 5 | 9 | 11 | 7 | 16 |
| 16 ID | 1.0000 | 5 | 4 | 9 | 7 | 6 | 14 | 18 | 7 | 2 | 16 |
| 9 MS | 1.0000 | 5 | 9 | 4 | 7 | 6 | 4 | 5 | 15 | 2 | 6 |
| 19 NB | 1.0000 | 5 | 9 | 4 | 7 | 6 | 2 | 4 | 16 | 18 | 6 |
| 24 AL | 1.0000 | 5 | 9 | 4 | 7 | 6 | 4 | 7 | 2 | 1 | 6 |
| 17 JM | 1.0086 | 9 | 5 | 4 | 15 | 6 | 4 | 11 | 13 | 16 | 6 |
| 20 PI | 1.0113 | 9 | 5 | 15 | 4 | 14 | 8 | 7 | 6 | 9 | 10 |
| 3 KN | 1.0390 | 9 | 15 | 14 | 2 | 5 | 18 | 16 | 6 | 10 | 17 |
| 10 AK | 1.0379 | 15 | 9 | 14 | 2 | 12 | 14 | 10 | 11 | 4 | 15 |
| 14 SE | 1.0311 | 15 | 9 | 14 | 2 | 11 | 15 | 12 | 9 | 18 | 14 |
| 4 SG | 1.1717 | 15 | 9 | 14 | 2 | 11 | 14 | 10 | 9 | 17 | 2 |
| 5 TB | 1.0762 | 15 | 2 | 14 | 11 | 9 | 2 | 15 | 4 | 17 | 5 |
| 23 SR | 1.4658 | 15 | 2 | 14 | 11 | 9 | 16 | 17 | 11 | 2 | 9 |
| 2 ES | 1.1644 | 15 | 11 | 2 | 14 | 13 | 15 | 9 | 8 | 11 | 14 |
| 1 EG | 1.1265 | 15 | 11 | 2 | 14 | 13 | 17 | 8 | 9 | 13 | 15 |
| 11 EB | 1.2101 | 15 | 11 | 2 | 13 | 14 | 14 | 3 | 15 | 12 | 6 |
| 13 CA | 1.6682 | 15 | 11 | 2 | 13 | 14 | 15 | 17 | 1 | 3 | 18 |

Ratio is the yield (or whatever the trait) for the winner within each environment (identified in the first column of AMMI1 ranks), divided by the yield for the overall winner, with both yields estimated by the AMMI1 model. Ratio automatically equals 1 for environments that are won by the overall genotype winner. This ratio assesses the importance of narrow adaptations, which are caused by GE interactions. A ratio of 1.05 or 1.10 means that narrow adaptations offer a yield increment of 5% or 10%. When the yield increment is large enough to have agricultural or economic significance, narrow adaptations are worthwhile, although at the cost of subdividing a growing region into two or more mega-environments.

In this table, mega-environments have different winners and they are separated by blank lines. Ordinarily small mega-environments are merged into adjacent larger mega-environments, especially when this imposes negligible loss in yield because an adjacent winner holds second (or third) rank in the merged environment. Often such mergers can simplify a mega-environment scheme considerably, producing a favorable tradeoff: a large gain in practicality, accompanied by a negligible loss of yield in only several of the environments.

When crafting a mega-environment scheme for a given yield trial, bear in mind that narrow adaptations caused by predictable GE interactions increase the number of usable mega-environments, whereas unpredictable GE interactions decrease this number. Often soil and management are predictable from year to year, whereas weather is unpredictable.

The fourth table in AMMISOFT output simply lists the ranked means and ranked IPC1 scores, first for genotypes and then for environments. It also provides the correlation between genotype means and IPC1 scores, and the correlation between environment means and IPC1 scores. These ranked lists should be inspected carefully to determine whether they have a plausible biological or ecological interpretation. If genotype means and IPC1 scores have a small correlation, then these means and scores call for different biological explanations; otherwise, if a large correlation, then the same explanation. The same principles apply to ecological interpretation of environment means and IPC1 scores. Genotype IPC1 scores and environment IPC1 scores require a coherent interpretation, such as drought tolerance for genotypes and rainfall for environments. If either genotypes or environments are better known than the other, begin with whichever is more familiar. Start by contrasting the top several genotypes (or environments) with the bottom several ones in order to suggest a biological (or ecological) interpretation, and then inspect the entire list to confirm a systematic trend and clear interpretation.

Finally, this particular wheat trial has no missing data, but if a dataset does have missing values (meaning zero replications for some genotype-environment combinations), then a fifth table lists the imputed values. AMMISOFT imputes missing values with the expectation maximization algorithm, specifically using AMMI1 rather than a higher member of this model family because AMMI1 tends to be most robust. At the bottom of this table, AMMISOFT gives the smallest and largest observations for the actual data, and the smallest and largest values for the imputed data. That information is helpful when scanning the imputed data to check for implausible values.

## FOUR GRAPHS

Four kinds of graphs have been especially frequent in the AMMI literature, and they can be produced by AMMISOFT. An explanation is available for each kind of graph about how to interpret it. Graphs can be produced in color (blue markers for genotypes and red markers for environments), or black-and-white. Graphs can be saved in a variety of formats, including jpg, pdf, png, ps, and tif.

The first kind of graph is the AMMI1 biplot. It has mean on the abscissa and IPC1 score on the ordinate, and it shows markers for both genotypes and environments, as in Fig. 1 in Gauch and Zobel (1997) and Fig. 1 in Gauch et al. (2008). Among markers of the same kind, displacements along the abscissa indicate differences in main (additive) effects, whereas displacements along the ordinate indicate differences in interaction (multiplicative) effects. IPC1 scores near zero indicate small interactions. A genotype and environment with IPC1 scores of the same sign have a positive GE interaction; whereas opposite signs mean negative interaction.

The second kind of graph is a linear model of genotype responses using the AMMI1 model, as in Fig. 2 in Gauch and Zobel (1997) and Fig. 3 in Gauch et al. (2008). It has environment IPC1 score on the abscissa and nominal yield (which ignores environment means) on the ordinate. This graph is analogous to the familiar Finlay and Wilkinson (1963) regressions that instead show genotype responses as linear functions of environment means. However, AMMI1 always captures as much or more of the GE interactions as do Finlay-Wilkinson regressions, often far more.

The third kind of graph is an AMMI1 biplot with horizontal lines added to display the mega-environment associated with each genotype winner, as in Fig. 3 in Gauch and Zobel (1997) and Fig. 2 in Gauch et al. (2008). Each winner occupies a horizontal band, and it wins all of the environments located within its band, and hence a horizontal band constitutes a mega-environment. The genotype winner for the mega-environment that includes the IPC1 score of zero is necessarily the yield trial’s overall winner.

The fourth and final kind of graph is an AMMI2 biplot showing IPC1 on the abscissa and IPC2 on the ordinate. When two or more IPCs are statistically significant, this graph can be of interest to display GE interaction patterns beyond those captured in an ordinary AMMI1 biplot. Markers near the origin have little GE interaction (at least presuming that IPC3 and higher axes are not significant or important). Two markers of the same kind (both genotypes or else both environments) in the same direction from the origin have similar interaction patterns; markers in opposite directions have opposite interactions; and markers nearly at right angles have uncorrelated interaction patterns. A genotype marker and an environment marker in the same direction from the origin and far from the origin have a large positive GE interaction; those in opposite directions have a large negative GE interaction. Only AMMI1 and AMMI2 biplots are common in the agricultural literature, but AMMISOFT computes up to 7 IPCs and any pair of IPCs can be selected for making a biplot. However, it is inappropriate to graph IPCs that lack statistical significance or some other evidence of agricultural relevance.

## CONCLUSION

The benefits from AMMI statistical analysis—when implemented with best practices—are considerable (Gauch, 2013). Yield can be increased by a practical mega-environment scheme that exploits narrow adaptations, based on the understanding of GE interactions that AMMI provides. Also, parsimonious AMMI models can gain accuracy, often as much as would doubling or tripling the number of replications, and this improves selections, increases reproducibility, and accelerates yield gains. Besides these advantages in conventional plant breeding, AMMI has applications in molecular breeding, such as scanning for quantitative trait loci (Gauch et al., 2011). AMMISOFT makes it easy for agricultural researchers to analyze their yield trials effectively.

## Supporting information

AMMISOFT documentation

Wheat yield trial

## SUPPLEMENTARY MATERIAL

AMMISOFTdocumentation.docx is the 17-page documentation for this software.

ESW8.txt is the wheat yield trial used in this preprint with 18 genotypes and 25 environments.

