## Supplementary material for "AMMISOFT for AMMI Analysis with Best Practices": AMMISOFT documentation

AMMI Analysis Using AMMISOFT

Hugh Gauch, Soil and Crop Sciences, Cornell University; September 2017.

**Download and Installation**. AMMISOFT is available for free from Hugh Gauch’s Cornell home page. The zipped download file uses about 44 MB, and the unzipped files about 134 MB of memory. While AMMISOFT is running, it requires about 256 MB of high-speed memory. AMMISOFT has been developed for PC computers with a Windows operating system. The user interface and graphs are coded in about 500 lines of Python and the AMMI computations and tables are coded in about 2,000 lines of FORTRAN. However, this software is self-contained so neither a Python interpreter nor a FORTRAN compiler are needed (unless users want to make modifications in their own version of this software).

A special feature of this software is that each of its four tables and four graphs are accompanied by explanatory material. The objective is to enable agricultural researchers to perform AMMI analysis with ease and yet with best practices.

**Program Execution**. The icon for AMMISOFT shows a green letter A flanked by tan heads of wheat; click on this icon to launch the program. Agree to the terms and conditions in order to open the main window and run AMMISOFT; alternatively you can open a window that shows the license terms. At the top of this main window click on “Choose file” which opens a browser to select your data. You can then click on “AMMI Analysis” which will report success if your dataset is alright, or else will show error messages. Following successful analysis, you can “View Tables” which will display the AMMI analysis in Notepad; or you can “Make Graphs.” The window for making graphs is very intuitive and simple: Four kinds of graphs are available, and you can choose whether to label genotypes or environments or both and can choose color or black-and-white. Graphs can be saved in a variety of formats, including jpeg and pdf. If you want to make additional graphs for a dataset that has already been analyzed, you can read its AMMISOFT output file and go directly to “Make Graphs” without repeating “AMMI Analysis.” Finally, at the bottom of the main window you can select “Data Format” for detailed instructions on the required format for datasets, or once you are familiar with that material, you can select “Data Format Short Guide” for a convenient one page summary.

**Page	Contents**

1	Installation and Program Execution

2–7	Data File Format for AMMISOFT

8–12	Four Tables

13–16	Four Graphs

17	Missing Data

**Data File Format for AMMISOFT**

**Introduction to AMMI Analysis and AMMISOFT Software.** The Additive Main effects and Multiplicative Interaction (AMMI) model uses analysis of variance for the additive or main effects followed by principal components analysis for the multiplicative or interaction effects. The data structure that AMMI addresses is a two-way factorial design, with or without replication. In agricultural research, the principal application is yield trials which have a number of genotypes tested in a number of environments, often with replication. Although yield is the foremost trait, others are also common, including plant height, days to flowering or maturity, disease resistance, and nutritional value. If there are multiple traits, AMMI analyses them individually, not all together. Also, when environmental factors such as rainfall or soil nitrogen have also been measured (beyond the yield trial itself), this additional information can help to interpret AMMI results, although this environmental data is not directly incorporated in AMMI analysis. When main and interaction effects are statistically significant and agriculturally important, as is most often the case, AMMI is extremely useful. The purpose of AMMISOFT is to make it easy to implement AMMI analysis with best practices as described in: Hugh G. Gauch, Jr., 2013, A Simple Protocol for AMMI Analysis of Yield Trials, Crop Science 53:1860–1869.

AMMISOFT uses the same format for datasets as did its predecessor MATMODEL (except for deletion of the optional format for single-entry lines of data). For all of the following items, enter full lines completing everything called for, even if this requires blanks or zeros at the end of a line. If the data were already entered into the computer for some other purpose, hopefully little work will be required to prepare the file for AMMISOFT. Data files must be created as plain text files and hence their filenames must end with .txt for the extension. Two or three lines precede the data, and they are supplied in the following order.

**Dataset Name.** The first line of the input data file identifies the dataset.  It has two entries.

(1)	Columns 1–72, an alphanumeric title for the dataset of up to 72 characters.  This title is reproduced once at the beginning of the AMMISOFT output in order to identify or describe your dataset.

(2)	Columns 77–80, an alphanumeric brief name for the dataset. This name is worth choosing carefully because it appears repeatedly in the AMMISOFT output file as a concise way to identify your dataset.  A useful convention is to use this brief name also in the dataset’s filename. For example, the supplied dataset with the brief name SOYB has been given the corresponding filename SOYB.txt. AMMISOFT automatically generates a name for its output file from the name for the input data file by adding the suffix “ammi” before the extension, so the result in this case is SOYBammi.txt. An output file is created in the same subdirectory as its corresponding input file so that these input and output files will appear together in your browser.

**Data Information.** The second line of the data file contains thirteen entries.

(1)	Columns 1–20, an alphanumeric name for the data’s measurement and units.  For example, “Yield  kg/ha” is the measurement and units for dataset SOYB.

(2)	Columns 21–25 right justified, a scaling integer.  This entry may be a positive or negative integer, or zero.  Ten is raised to the power of this integer, and then the data are multiplied by the result.  If this entry is 0 (or blank), then no action is taken.  Scaling can be useful if the data have been entered already as rather small or large numbers, or to avoid having to enter decimal points.  The measurement units given in the previous entry should reflect the result after multiplication by the indicated scaling factor.

*In order to produce results which print nicely in the print formats used by AMMISOFT, it is required that typical data entries be in the range of 0.001 to 1000000, or better yet within 0.01 to 100000, or ideally within 1 to 10000.* If necessary, the units of measurement should be changed in order to result in values in this range.

(3)	Columns 26–30 right justified, number of genotypes.  For data file SOYB, there are 7 genotypes.

(4)	Columns 32–35 left justified with 1 to 4 characters, brief name for genotypes.  As explained soon, three such names are given:  one for genotypes, environments, and replications.  The first letter of each of these three names must be unique (in columns 32, 42, and 52).  For SOYB, the name “GEN” is given for the genotypes.

(5)	Columns 36–40 right justified, number of environments. For SOYB, there are 10 environments.

(6)	Columns 42–45 left justified with 1 to 4 characters, brief name for environments.  Remember that the first letter of this name in column 42 must be unique (not identical to the entry in column 32 or 52).  For SOYB, the name “ENV” is given for the environments.

(7)	Columns 46–50 right justified, number of replications.  If the data are unreplicated, then this entry should be 1. If the data are unbalanced, with various treatments having various numbers of replicates, then this entry should be the largest number of replicates encountered. For example, if various treatments have 2 to 4 replicates, then this entry should be 4.

(8)	Columns 52–55 left justified with 1 to 4 columns, brief name for replications.  Remember that the first letter of this name in column 52 must be unique (not identical to the entry in column 32 or 42).  For SOYB, this and the previous entry indicate 4 REP for replications.

(9)	Columns 58–60, order of data entries. Subscripts for genotypes, environments, and replicates are identified by the first letters of their brief names, which are located in columns 32, 42, and 52.  The order of data entries is specified by listing these subscripts according to their rate of change, *the first changing fastest and the last changing slowest.* If only one replicate is read, replication changes most slowly (in fact, never), so it is listed last (in the third position; it is *not* omitted).  For the SOYB data, the first data entry 2729 is for genotype 1, environment 1, and replicate 1.  The second entry, 2662, is for genotype 2, environment 1, and replicate 1.  Hence, genotype changes fastest, so it is listed first.  There are 7 genotypes.  The eighth data entry, 2747, is for genotype 1, environment 1, and replicate 2, so replicates change second fastest.  There are 7x4 = 28 entries for the first environment, and the twenty‑ninth entry, 983, begins the data for environment 2, the subscript for environment thus changing slowest.  Consequently, the data order specification for this dataset is GRE. Note that the subscript code letters given earlier in columns 32, 42, and 52, and the letters given here in columns 58–60 must match exactly, including case.

(10)	Columns 62–65 left justified with 1 to 4 characters, brief name for treatments. A treatment is defined as a genotype-and-environment combination. The first letter of this name is *not* required to be unique relative to the first letters of the names for genotypes, environments, and replications (in columns 32, 42, and 52). For SOYB, this brief name is “TRT.”

(11)	Columns 68–70 left justified with 1 to 3 characters, experimental design. For SOYB the experimental design is RCB for the randomized complete block design. Likewise, LAT could indicate a lattice design, or RAN a completely randomized design. The one and only experimental design that AMMISOFT is designed to recognize and analyze is the randomized complete block design, denoted by RCB, and this design makes possible calculation of an error mean square to be used for significance tests.

(12)	Columns 71–75 right justified or else including decimal point, missing value indicator. This value must be numeric. Data entries with this value will be marked by AMMISOFT as missing data. Provided that no actual observations happen to have a value of zero, the missing value indicator may be set to zero in order to mark entries of zero as missing observations. A zero may be entered by blanks in columns 71–75, but it is better to enter an explicit “0” or “0.0” in order to make this entry visible. Note that a missing value indicator of zero causes data entries of zero to be treated as missing values, not as actual observations of zero. If there are no missing data, and observations are always positive, then a specification of zero will have the intended effect of marking no data as missing. Likewise, if there are no missing data, but some observations are zero although none are negative, then a missing value indicator of –1 will have the intended effect of marking no data as missing.

(13)	Column 80, format supplied or not. If column 80 contains “N” (in upper or lower case), then no format is supplied and AMMISOFT reads the data in free format. Free format is suitable if no extraneous information is interspersed with the data, if a line break or at least one space always separates entries, and if entries of zero are given as “0” or equivalent rather than as a blank. If these conditions are not met, enter “F” (in upper or lower case) in column 80 and also supply the format, as explained momentarily.

If no format is supplied, that is if the second line contains “N” in column 80, then the following data lines are to be read in free format. Given G genotypes, E environments, and R replications, exactly GER entries must follow. Missing values, if any, are indicated by the missing value indicator (specified in columns 71–75 of line 2). Free format allows data entries to appear anywhere on a line, but all entries must be separated by a line break or tab or at least one space, and there must be no extraneous information, including no commas.

**Optional Data Format.** Another line of the input data file is optional, included only if the format is supplied, that is, if column 80 of the second input line contains “F” rather than “N.” AMMISOFT uses extremely fast code written in FORTRAN to read the data file and to perform AMMI analysis (and additional code written in Python to read the output file and to make graphs). This format line contains one item, the supplied FORTRAN format in columns 1–72. Be sure to include the enclosing left and right parentheses. Field specifications for data entries must be real, not integer, because these entries will be read as single‑precision real numbers. Even if the data entries are typed as integers that are right justified in their fields, they must still be read as real numbers.

The option F is taken (in column 80 of line 2) when the data are to be read with a supplied format that uses the FORTRAN rules for a format statement. A supplied format is needed if the data lines contain extraneous information interspersed with the data entries, if line breaks or spaces do not always separate entries, or if entries of zero are indicated by blanks. Three examples follow of valid format specifications for this option F:

(7F9.2)

(5F5.0, 10X, F5.0, 15X, 2F5.0)

(3(10F8.3), /, 4F8.3)

The first and second examples are rather simple. The third applies to data organized in sets of four lines, with three lines similar and the fourth different.

Explicit decimal points in data entries override the format statement. For example, the format specifier F5.0 will correctly read all of the following fields of 5 characters where “^” is used to indicate the presence of a blank column: ^2.3^, ^1653, 1653., 1.2E3, and ^^^^5. However, 1653^ would *not* be read as 1653, but rather as 16530. Values entered like integers, without decimal points, must be right justified in their fields in order to be read correctly.

**Data Entries.** The data entries then follow in the input data file. Values may be positive, negative, or zero, but values equaling the missing value indicator will be marked as missing data. The data are stored in single precision, so up to about 7 significant digits can be retained. For a specified format, be sure to enter complete data lines satisfying the entire format, even if some entries are zero (regardless whether a zero represents an actual observation of zero or else a missing observation). For example, in option F with format (4F5.0), given a line containing only two actual observations of 2541 and 2619 and two missing observations indicated by values of zero, this line must still contain the full 20 characters called for by this format, although two of the fields (of 5 characters each) may be filled with blanks.

As noted earlier regarding scaling, in order to produce results which print nicely in the print formats used by AMMISOFT, it is required that typical data entries be in the range of 0.001 to 1000000, or better yet within 0.01 to 100000, or ideally within 1 to 10000.

Because the order of change of subscripts can be specified, if the data have already been entered into the computer for other purposes, the data may already be nearly in a suitable format for AMMISOFT.  But if the data file contains any extraneous information, such as line numbers or code names for anything, a specified format should be used in order to ignore this extraneous information.  In unusually complex cases, a small computer program may need to be written to transcribe the data into a format acceptable to AMMISOFT.

**Genotype and Environment Code Names.** Next the input data file contains 4‑character code or brief names for the genotypes and environments.  The format is 4 characters followed by a blank, with 15 entries per line.  The names use alphanumeric characters and may contain blanks.  Genotype names are given first, and then, beginning on a new line, environment names.  For SOYB, the 7 genotype code names use one line, and the 10 environment code names (indicating location-and-year combinations) occur on the following line.

Alternatively, these names can be omitted by ending the data file at the last data entry, with no spaces or anything else following. Then sequential integers are substituted for these names.

**Example.** The following simple example illustrates a correct format to be read by AMMISOFT. It is the supplied data file TOYD.txt that appears on pages 55 and 86 of my text on AMMI: Hugh G. Gauch, Jr., 1992, Statistical Analysis of Regional Yield Trials: AMMI Analysis of Factorial Designs, Elsevier.

A ruler has been placed above this example to facilitate determining column positions.

```
         1         2         3         4         5         6         7         8
12345678901234567890123456789012345678901234567890123456789012345678901234567890

Toy data, Table 3.1 page 55 in Gauch 1992 Elsevier text on AMMI.            TOYD
Yield  kg/ha            0    5 GEN     4 ENV     3 REP   RGE TRT   RAN    0    N
 352 298 331   249 295 251   266 226 219   231 254 196   182 199 216
 268 271 232   253 270 218   243 215 205   199 205 217   234 184 221
 201 170 169   165 214 206   154 206 153   157 176 144   171 159 195
 125 121 102   180 133 158   159 146 124   170 180 139   121 138 164
GEN1 GEN2 GEN3 GEN4 GEN5
ENV1 ENV2 ENV3 ENV4
```

**Two Additional Examples**. Dataset SOYB.txt has data from a New York soybean trial that appears on page 56 of my text. Dataset ESW8.txt has an international wheat trial published in Crossa, Fox, Pfeiffer, Rajaram, and Gauch (1991), Theoretical and Applied Genetics 81:27–37. It was also used as the main example in my 2013 paper on AMMI protocol that was already cited.

Supplied dataset SOYB.txt is shown below with data for 10 soybean genotypes, 7 New York environments (location and year combinations), 4 replications, and hence 280 observations. The experimental design is randomized complete blocks (RCB). These data were published on page 56 in: Hugh G. Gauch, Jr., 1992, Statistical Analysis of Regional Yield Trials: AMMI Analysis of Factorial Designs, Elsevier.

The AMMISOFT analysis follows, first four tables and then four graphs. Each of these eight items has an accompanying explanation of its mathematical and agricultural meaning.

```
New York soybean yields, Table 3.2 in HGG text, page 56.                    SOYB
Yield  kg/ha            0    7 GEN    10 ENV     4 REP   GRE TRT   RCB  0.0    N
  2729  2662  2638  2680  2598  2908  2732
  2747  2238  2425  2072  3036  3430  2951
  2593  2454  2191  2994  3097  3265  2432
  2832  2528  2079  3218  2641  2823  2866
   983   824  1286  1299  2196  2159  1529
  1445   491  1258  1484  2239  2041  1767
  1013   381  1390  1646  1932  1463  2079
  1004   616  1178  1568  1486   981  1472
  1525  1484  2597  2404  2798  2170  2903
  2348  1426  2686  3028  2500  3125  2944
  1968  1091  2198  2742  2520  2825  2428
  2309  1545  1920  3028  2650  3750  2757
  1783  1902  1717  1707  1419  1814  1378
  1957  1534  1405  1955  1404  1967  1656
  1609  1416  1591  1825  1414  1615  1508
  1593  1574  1637  1870  1262  1797  1593
  3194  2619  2742  3561  2760  3039  2285
  3085  2839  2560  3208  2911  3250  3119
  3489  3263  2742  3213  3569  3275  3305
  3263  3121  3207  3709  3589  3207  2992
  2989  2652  3317  3764  2900  3639  3715
  3640  2366  2334  2862  3298  3647  3641
  3105  2835  2815  3276  3188  3613  3353
  2922  2570  2794  3727  3367  3677  3650
  2574  1989  2602  2892  2219  2840  2726
  2312  2394  2322  2368  2674  2467  2363
  2495  2138  2443  2941  2786  3067  2580
  2793  2617  2752  3122  2655  2438  2484
  2961  3348  2206  2484  1267  2177  1504
  3354  2909  2313  2467  1220  1324  1630
  3470  3169  2357  2302  1257  1998  1577
  3307  3139  2429  3041  1207  1363  1932
  4370  3852  3185  2824  2022  2770  3192
  3726  3605  3300  3361  2920  3503  2783
  3818  3532  3042  3362  2353  2666  3005
  3780  3461  2861  3111  2633  2350  2626
  5437  4687  3085  4275  4615  4777  4447
  5165  4650  4372  4632  4756  4941  3509
  4273  4749  3492  5522  4243  3790  2750
  4750  4347  3090  4578  4392  3328  4053
EVAN WILK CHIP HODG S200 CORS WELL
A77  V79  R81  I85  G85  A86  N87  C87  C88  G88
```

First Table (prefaced with basic information about the dataset)

```
AMMISOFT for SOYB:  Data Input

New York soybean yields, Table 3.2 in HGG text, page 56.

Data input filename:
C:/hg/AMMIforSOYB/SOYB.txt

AMMI output filename:
C:/hg/AMMIforSOYB/SOYBammi.txt

Measurement  Yield  kg/ha
Number of GEN       7
Number of ENV      10
Number of REP       4

Number of TRT  with data        70 out of        70 possible.
No empty cells.

Number of observations         280 out of       280 possible.
No missing data; these data are balanced.

Required memory is     .0010% of allocated memory.

Grand mean     2678.19643


AMMISOFT for SOYB:  ANOVA Table

-------------------------------------------------------------------------------
Source            df               SS                  MS          Probability
-------------------------------------------------------------------------------
Total             279       244450370.19643        876166.20142
  TRT              69       223206475.94643       3234876.46299    .0000000 ***
    GEN             6         7117668.77143       1186278.12857    .0000000 ***
    ENV             9       176360099.37500      19595566.59722    .0000000 ***
    GxE            54        39728707.80000        735716.81111    .0000000 ***
      IPC1         14        32756258.12665       2339732.72333    .0000000 ***
      IPC2         12         4681150.07194        390095.83933    .0033084 **
      IPC3         10         1019856.87543        101985.68754    .6528434
      IPC4          8          801796.60736        100224.57592    .7515836
      IPC5          6          430131.36477         71688.56079    .8877453
      Residual      4           39514.75386          9878.68846    .9805159
  Error           210        21243894.25000        101161.40119
    Blocks/Env     30         4372037.96429        145734.59881    .0420758 *
    Pure Error    180        16871856.28571         93732.53492
-------------------------------------------------------------------------------
F-tests use Pure Error because Blocks/Env are significant at the 0.05 level.

MODEL DIAGNOSIS:

AMMI comprises a model family with AMMI0, AMMI1, AMMI2 and so on
retaining 0, 1, 2, or more interaction principal components (IPCs)
before relegating higher components to a discarded residual;
finally, the full model AMMIF equals the actual data (or averages
over reps), so it has no residual.  Model diagnosis is required
to determine the best AMMI model for a given dataset, based on
statistical and practical considerations.

FR-tests at the 0.01 level diagnose AMMI2.

Estimated sums of squares for GxE signal and noise:
GxE total           39728707.80000
GxE noise            5061556.88571 or  12.74%
GxE signal          34667150.91429 or  87.26%
Early IPCs selectively capture signal, and late ones noise.
Accordingly, this much signal suggests AMMI1 or maybe AMMI2.

Conclusion:
The consensus diagnosis is AMMI2 for this dataset.

Perspective:
Note that the SS for GE-signal is  4.87 times that for GEN  main effects.
Hence, narrow adaptations are important for this dataset.
Even just IPC1 alone is  4.60 times the GEN  main effects.
Also note that GE-noise is   .71 times the GEN  main effects.
Discarding noise improves accuracy, increases repeatability,
simplifies conclusions, and accelerates progress.
```

Second Table

```
AMMISOFT for SOYB:  Winners for AMMI Model Family

-------------------------------------------------
              AMMI Model Family
              -----------------------------------
Genotype          0    1    2    3    4    5    F
-------------------------------------------------
    1  EVAN            4    4    4    3    3    3
    4  HODG      10    2    3    2    3    3    3
    6  CORS            4    3    3    3    3    3
    5  S200                      1    1    1    1
-------------------------------------------------
Mega-environments 1    3    3    4    4    4    4
-------------------------------------------------
Another    3 genotypes never win, so they are not listed.

     Mega-environments are distinguished by having different genotype
winners.  Increasingly complex AMMI models generally have more genotype
winners or m-es, as shown in the list at the bottom of the above table.
This is an important reason why model diagnosis matters.
     Mega-environment analysis has been applied mostly to yield-trial
data, for which larger values are better.  It is also applicable to
other traits, such as disease resistance, but again larger values must
indicate better genotypes.  For instance, if the original scale is
0 for most resistant to 5 for most diseased, then subtract those values
from 5 in order to obtain a transformed scale with 0 worst and 5 best.
     In the above table, the genotypes are listed in IPC1 order,
so those at the top and bottom have opposite GxE interaction patterns.
Often this contrast in the genotypes has an evident agricultural
interpretation that has a corresponding contrast in the environments
(which are listed in their IPC1 order in the following table).
A genotype at the top of this table has positive GxE interactions with
environments at the top of the next table and negative GxE interactions
with environments at the bottom of the next table; and the opposite
patterns apply to genotypes at the bottom of this table.
     AMMI models to the left of the best AMMI model (for optimizing
predictive accuracy) are too simple, so they underfit real signal;
whereas models to the right of the best one are too complex, so they
overfit spurious noise.  At the opposite extremes, AMMI0 always captures
no GE-signal and no GE-noise, whereas AMMIF always captures all GE-signal
but with this also all GE-noise.  Accordingly, a parsimonious intermediate
AMMI model is often most predictively accurate, such as AMMI1 or AMMI2,
because missing little GE-signal while discarding much GE-noise
is better than either of the opposite and worse problems of missing a
huge amount of GE-signal or capturing a huge amount of GE-noise.
This opportunity to increase predictive accuracy is another reason why
model diagnosis matters.
     Practical constraints frequently limit the number of workable m-es
to only 2 or 3 (or perhaps a few more).  This can require a lower AMMI
model than that diagnosed solely by statistical considerations, often
AMMI1, in order to achieve fewer m-es.  Often this lower AMMI model is
almost as predictively accurate as the best model, and is also far more
accurate than AMMIF, that is, the actual data (or averages over reps).
     The above table displays the genotype winners and the consequent
numbers of m-es for a wide spectrum of options from AMMI0 to AMMIF.
Thereby it illuminates the tradeoff between statistical and practical
considerations when choosing the most appropriate member of the AMMI
model family to use for a given dataset.
```

Third Table

```
AMMISOFT for SOYB:  Ranking Table for AMMI1 and AMMIF

----------------------------------------------------------------------------
                              AMMI1 Ranks                  AMMIF Ranks
                      -------------------------    -------------------------
Environment   Ratio       1    2    3    4    5        1    2    3    4    5
----------------------------------------------------------------------------
    8  C87   1.3208       1    2    4    3    6        1    2    4    3    6
    9  C88   1.1410       1    2    4    3    6        1    2    4    3    7
   10  G88   1.0528       1    4    2    6    3        1    4    2    5    6
    4  I85   1.0066       1    4    6    2    7        4    6    1    2    3

    5  G85   1.0000       4    1    6    7    5        4    1    5    6    2
    7  N87   1.0000       4    6    1    5    7        4    6    5    1    7

    1  A77   1.0029       6    4    5    7    1        6    5    7    4    1
    6  A86   1.0350       6    4    5    7    1        6    7    4    5    1
    2  V79   1.1619       6    5    7    4    3        5    7    6    4    3
    3  R81   1.1162       6    5    7    4    3        6    4    7    5    3
----------------------------------------------------------------------------
Switch from GEN      1 to     4 at ENV  IPC1 score           1.4282632
Switch from GEN      4 to     6 at ENV  IPC1 score          -6.3958343

     In the above table, the environments are listed in IPC1 order,
so those at the top and bottom have opposite GxE interaction patterns.
     Ratio is the yield (or whatever the trait) for the winner within
each environment (identified in the first column of AMMI1 ranks),
divided by the yield for the overall winner (which is genotype     4),
with both yields estimated by the AMMI1 model.  Ratio automatically
equals 1 for the overall winner.  This ratio assesses the importance
of narrow adaptations, which are caused by GxE interactions.  When a
5% or 10% yield increment has agricultural or economic significance,
a ratio of 1.05 or 1.10 or more indicates that narrow adaptations offer
substantial opportunities for yield increases, although at the cost of
subdividing a growing region into two or more mega-environments.
     M-es have different winners and they are separated by blank lines.
Ordinarily small m-es are merged into adjacent larger m-es, especially
when this imposes negligible loss in yield because an adjacent winner
holds second (or third) rank in the merged environment.  Often slight
editing of a ranking table, obtained by deleting or moving some blank
lines, can simplify the m-e scheme considerably.  Editing may achieve
a favorable tradeoff:  a large gain in practicality, accompanied by a
negligible loss of yield in several of the environments.
     At the bottom of the above table, the environment IPC1 scores are
listed for each switch from one winning genotype to the next.  The
following tables list IPC1 scores for all genotypes and environments.
     When crafting a m-e scheme for a given dataset, bear in mind that
narrow adaptations caused by predictable GxE interactions increase the
number of usable m-es, whereas unpredictable GxE interactions decrease
this number.  Often soil and management are predictable from year to
year, whereas weather is unpredictable.
```

Fourth Table (followed by computer-readable parameters)

```
AMMISOFT for SOYB:  Ranked Means and IPC1 Scores

------------------------------------------------------------------
 GEN   Code          Mean        |  GEN   Code      IPC1 Score
------------------------------------------------------------------
    4  HODG          2903.05000  |     2  WILK          32.4830948
    1  EVAN          2867.75000  |     1  EVAN          25.0464617
    6  CORS          2781.97500  |     3  CHIP           1.2918357
    7  WELL          2605.40000  |     4  HODG            .3311285
    5  S200          2599.82500  |     7  WELL         -18.1339352
    2  WILK          2525.42500  |     6  CORS         -18.5991618
    3  CHIP          2463.95000  |     5  S200         -22.4194237
------------------------------------------------------------------
Grand mean           2678.19643
Correlation between GEN  means and IPC1 scores   .0323401

------------------------------------------------------------------
 ENV   Code          Mean        |  ENV   Code      IPC1 Score
------------------------------------------------------------------
   10  G88           4310.89286  |     8  C87           34.0256597
    6  A86           3202.00000  |     9  C88           20.6809622
    9  C88           3143.32143  |    10  G88           11.1255632
    5  G85           3111.28571  |     4  I85            1.9227689
    1  A77           2709.25000  |     5  G85            -.8858324
    7  N87           2573.32143  |     7  N87           -4.4847079
    3  R81           2416.75000  |     1  A77           -6.8490192
    8  C87           2275.42857  |     6  A86          -12.7288471
    4  I85           1639.35714  |     2  V79          -20.2411462
    2  V79           1400.35714  |     3  R81          -22.5654012
------------------------------------------------------------------
Grand mean           2678.19643
Correlation between ENV  means and IPC1 scores   .2759915

     The above ranked lists should be inspected carefully to determine
whether they have a plausible biological or ecological interpretation.
If genotype means and IPC1 scores have a small correlation, then these
means and scores call for different biological explanations; otherwise,
if a large correlation, then the same explanation.  The same principles
apply to ecological interpretation of environment means and IPC1 scores.
     Genotype IPC1 scores and environment IPC1 scores require a coherent
interpretation, such as drought tolerance for genotypes and rainfall for
environments.  If either genotypes or environments are better known than
the other, begin with whichever is more familiar.  Start by contrasting
the top several genotypes (or environments) with the bottom several ones
to suggest a biological (or ecological) interpretation, and then inspect
the entire list to confirm a systematic trend and clear interpretation.


AMMISOFT for SOYB:  Computer-readable AMMI Parameters
    7   10    4    5          2678.19643
    1  EVAN          2867.75000          25.0464617           3.7808024          -5.2391265           1.5644712           9.7797608
    2  WILK          2525.42500          32.4830948           4.8035717          -2.4564616           4.2016569          -8.7391110
    3  CHIP          2463.95000           1.2918357         -21.3952614          12.9189971           3.1739461          -2.7408806
    4  HODG          2903.05000            .3311285           5.1654618           6.9192676         -16.5374806           3.8546439
    5  S200          2599.82500         -22.4194237          20.1018900           7.7012664           9.1944170            .5245229
    6  CORS          2781.97500         -18.5991618            .0296654         -10.4118083          -6.1947664          -9.3915498
    7  WELL          2605.40000         -18.1339352         -12.4861300          -9.4321346           4.5977557           6.7126138
    1  A77           2709.25000          -6.8490192           3.9826427          -9.4140015           5.9855932          -9.6410578
    2  V79           1400.35714         -20.2411462            .4210351           8.2182435          12.2917937           5.7180629
    3  R81           2416.75000         -22.5654012          -7.6369375           3.4267931         -11.2548838           3.1741578
    4  I85           1639.35714           1.9227689          -6.2052432            .8345696          -2.0519776         -10.1802321
    5  G85           3111.28571           -.8858324           6.4847006           4.7020417          -1.2025960          -1.0154815
    6  A86           3202.00000         -12.7288471          -3.5470256         -14.7614442          -4.0807815           6.0900561
    7  N87           2573.32143          -4.4847079          -3.3800750           7.8629845           -.1891044          -3.7883247
    8  C87           2275.42857          34.0256597          -9.1354529           3.6226094          -4.5248165           1.9789747
    9  C88           3143.32143          20.6809622          -8.3608830          -4.6218272           8.7853071           5.0169063
   10  G88           4310.89286          11.1255632          27.3772388            .1300311          -3.7585343           2.6469384
```

First Graph: AMMI1 Biplot

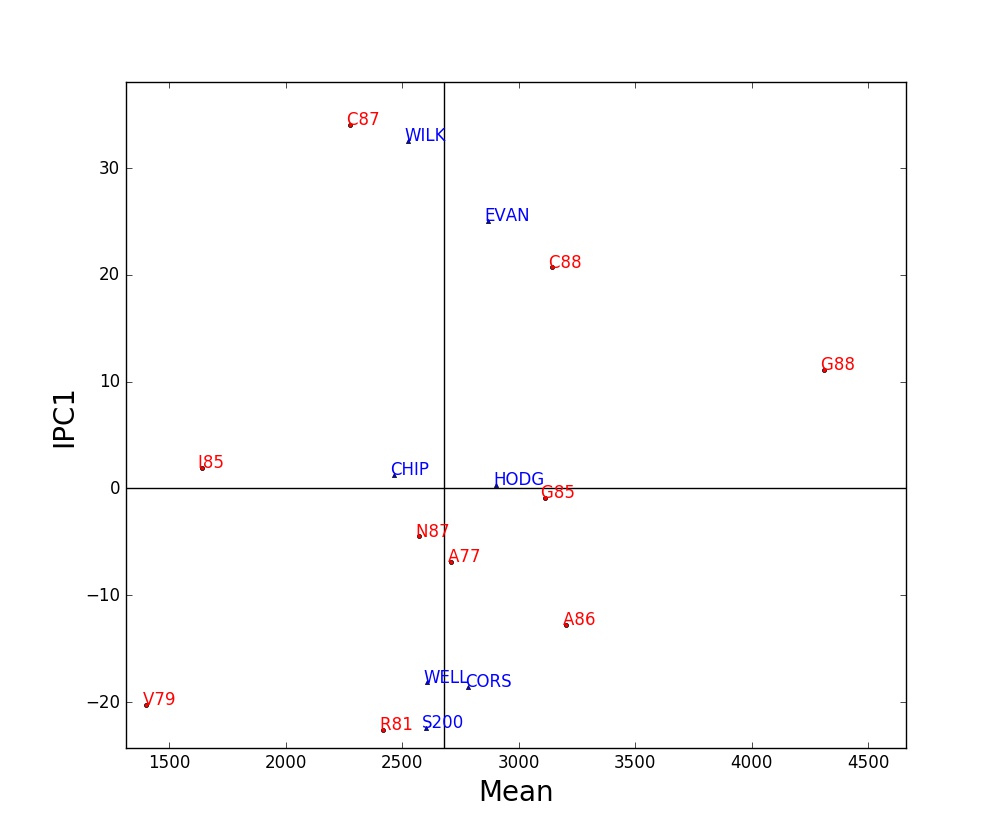

An AMMI1 biplot shows main and interaction effects for both genotypes and environments. Among markers of the same kind, displacements along the abscissa indicate differences in main (additive) effects, whereas displacements along the ordinate indicate differences in interaction (multiplicative) effects. IPC1 scores near zero indicate small interactions. A genotype and environment with IPC1 scores of the same sign have a positive GxE interaction; whereas opposite signs mean a negative interaction.

The legend for an AMMI1 biplot should report the percentage of the treatment sum of squares captured by genotypes, environments, and IPC1. Often they total over 90%. Sometimes this biplot captures virtually all of the variation that is signal rather than spurious noise – or at least it captures as much GxE interaction as is practical in terms of staying within a manageable number of genotype winners and hence mega-environments.

Attempt to give IPC1 scores a plausible biological, ecological, or geographical interpretation by contrasting genotypes at opposite extremes – and do the same for environments, seeking a coherent story. Often IPC1 is interpretable as a gradient in temperature or rainfall or such, but only rarely is IPC2 interpretable even when statistically significant.

Second Graph: AMMI Linear Model

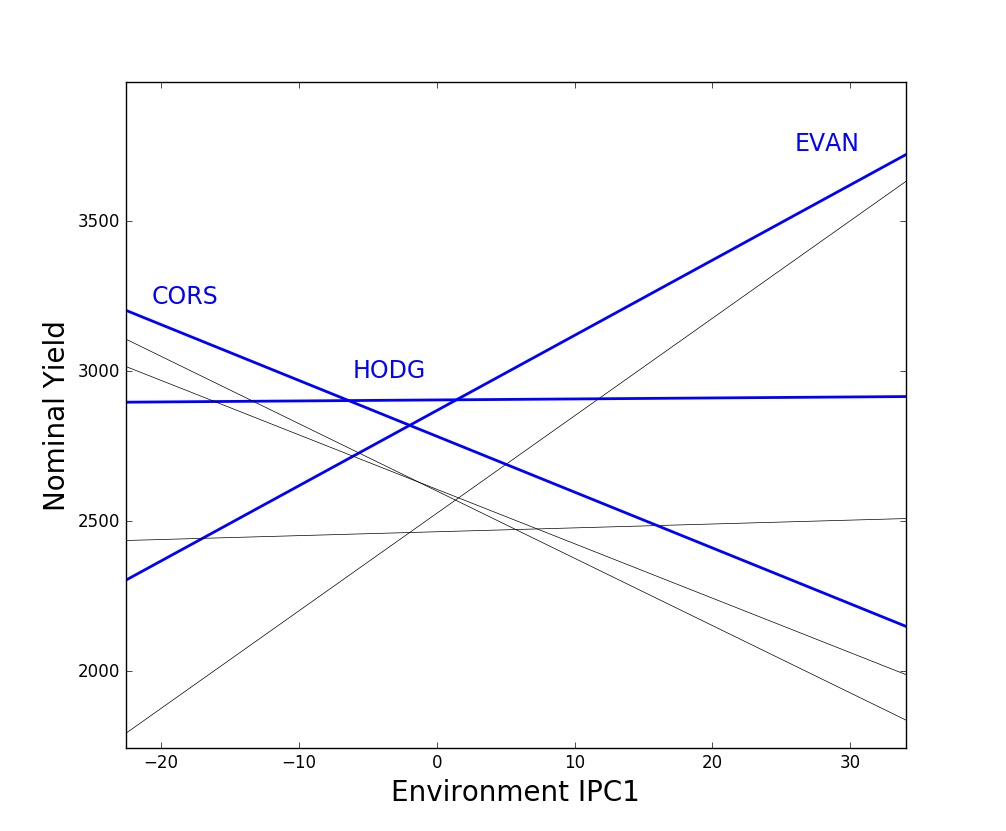

For the AMMI1 model, genotype responses are linear functions of environment IPC1 scores. This is analogous to the familiar Finlay-Wilkinson regressions that show genotype responses as linear functions of environment means. However, AMMI1 always captures as much or more of the GxE interactions as F-W regressions, often far more.

Genotype responses are shown by nominal yields that equal the genotype mean plus the AMMI1 estimated interaction (genotype IPC1 score times environment IPC1 score). But nominal yields exclude environment deviations from the grand mean because they are irrelevant for genotype rankings in a given environment, and adding them would make this graph quite messy.

Of special interest are the genotypes that win, which are at the top of this stack of lines for at least a portion of the range of environment IPC1 scores. This graph based on AMMI1 often has a small and manageable number of genotype winners, such as 2 to 5 winners – and any genotypes that win only tiny regions can be omitted with negligible loss of yield in order to achieve fewer winners and hence manageable mega-environments. By contrast, AMMI2 and higher models often implicate an impractical multitude of winners. Also, IPC1 usually has an evident and useful agroecological interpretation for both genotypes and environments, whereas higher IPCs are rarely interpretable.

Third Graph: AMMI1 Winners

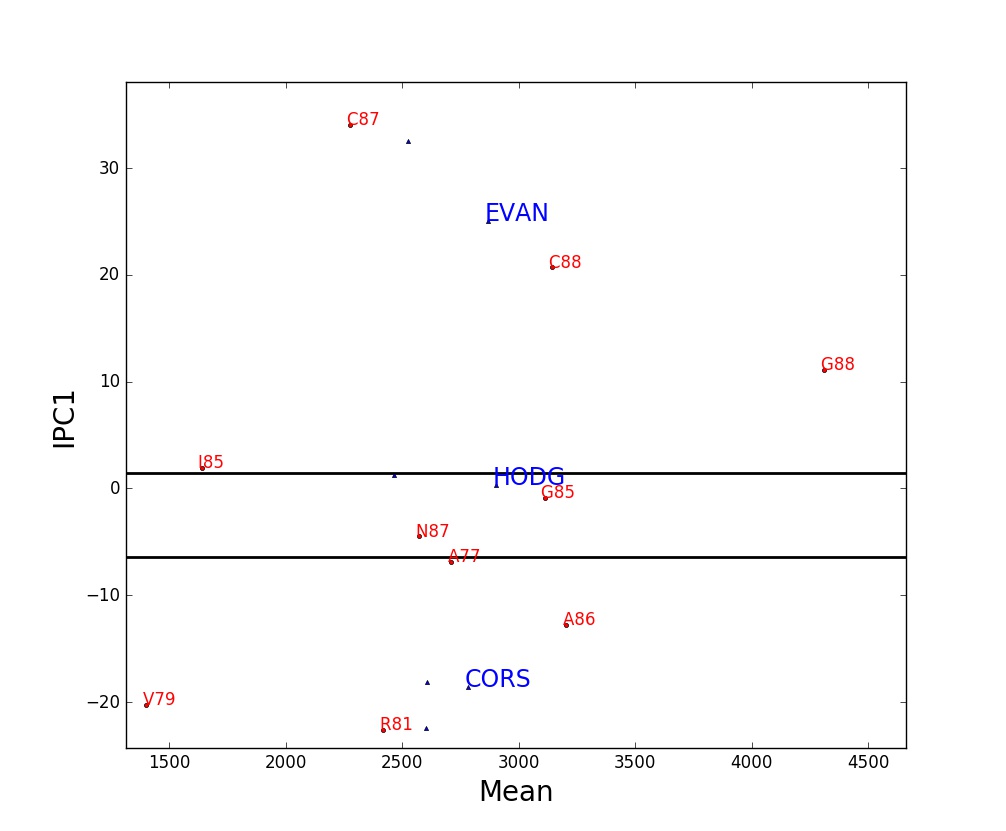

This AMMI1 biplot has been annotated to display the mega-environment associated with each genotype winner. Each winner occupies a horizontal band, it wins in all environments located within its band, and hence a horizontal band constitutes a mega-environment. Mega-environments are separated by horizontal lines, which means that they are determined by IPC1 scores solely. The genotype winner for the mega-environment that includes the IPC1 score of zero is necessarily the yield trial’s overall winner.

Attention is drawn to the genotype winners by writing their code names with a large font. From top to bottom, the order of the winners and of the mega-environments always corresponds – for instance, the genotype winner highest in this graph is the one that wins in the top mega-environment. However, the code names for winners are not necessarily written within their corresponding mega-environments.

An AMMI1 mega-environment display has a simple geometry and straightforward interpretation. The IPC1 scores and the consequent sequence of mega-environments often have a plausible and coherent interpretation as an environmental gradient in rainfall or whatever, along with cultivar variation in drought resistance or whatever. When the mega-environment for the overall winner contains rather few of the environments, specific adaptations are likely to offer substantial opportunities.

Fourth Graph: AMMI2 Biplot

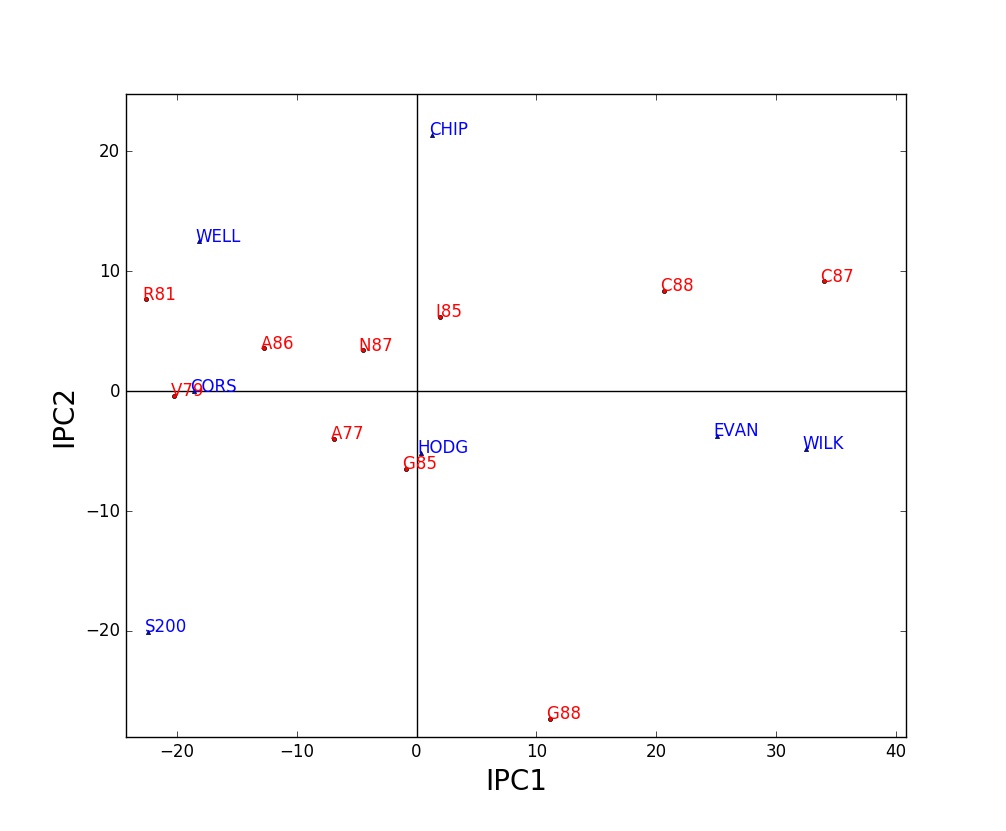

When two or more IPCs are statistically significant, they can be graphed to display GxE interaction patterns beyond those captured in an ordinary AMMI1 biplot. By far the most common such biplot shows IPC1 on the x-axis and IPC2 on the y-axis, and this is termed an AMMI2 biplot.

In an AMMI2 biplot, markers near the origin have little GxE interaction (presuming that IPC3 and higher axes are not significant or important). Two markers of the same kind (both genotypes or else both environments) in the same direction from the origin have similar interaction patterns; markers in opposite directions have opposite interaction patterns; and markers nearly at right angles have uncorrelated interaction patterns. A genotype marker and an environment marker in the same direction from the origin have a large positive GxE interaction; those in opposite directions have a large negative GxE interaction. More specifically and more generally, the AMMI2 model estimates the interaction for a given genotype in a given environment by the product of their IPC1 scores plus the product of their IPC2 scores.

Only AMMI1 and AMMI2 biplots are common in the agricultural literature, but AMMISOFT computes up to seven IPCs and any pair of IPCs can be selected for making a biplot. However, do not use IPCs that lack statistical significance or some other evidence of agricultural relevance.

**Missing Data**. If a dataset has missing values, which means no replicates for some genotype-and-environment combinations, then a list of values imputed by the expectation maximization model EM-AMMI1 is added to the AMMISOFT output. This list appears after the fourth table and before the computer-readable AMMI parameters.

For instance, the soybean dataset SOYB can be made incomplete by removing the data for three of its entries: EVAN in A77, WILK in G85, and CORS in C87. That is, for each of these entries, its four replicates of actual data are all changed to zeroes since zero is the missing data indicator for this dataset. This results in the following output:

```
AMMISOFT for SOY3:  List of EM-AMMI1 Imputed Values

----------------------------------
 ENV     GEN         Imputed Value
----------------------------------
    1       1           2737.39978
    5       2           2888.73133
    8       6           1801.75701
----------------------------------
Grand mean              2678.57697

Actual data              578.00000 to          4906.25000
Imputed values          1801.75701 to          2888.73133

     This dataset has  4.286% empty cells with no data.  Inspect these
imputed values to check for any that are especially implausible.
```

For comparison, the actual data (averages over four replicates) are 2725.25, 2960.50, and 1715.50 kg/ha. Hence, the imputing errors are 12.14978, –71.76867, and 86.25701. Their root mean square is 65.16296, or about 2.4% of the grand mean. This accuracy is satisfactory.

When a dataset has actual missing data, imputation accuracy can still be checked by generating additional missing data by temporarily withholding some data, and then comparing these actual and imputed values. This expedient works best when the distribution of the withheld missing data in the data matrix is made similar to the distribution of the actual missing data.

The EM-AMMI1 model is used to impute missing data, rather than a higher member of the EM-AMMI model family, because it tends to be most reliable. After imputing missing data, AMMISOFT then fits the ordinary AMMI model with 7 components (or fewer for a small dataset).

Extensive experience has shown that EM-AMMI1 generally works quite well. Nevertheless, missing data are still missing data. Exercise caution when interpreting results, especially if there is a lot of missing data, or the dataset is small (fewer than 10 genotypes or environments), or missing data are concentrated in certain rows or columns of the data matrix rather than missing data being fairly evenly scattered throughout the matrix.
